# COPD Preferentially Disrupts Constitutive Redox Support Relative to Inducible Antioxidant Defense

**DOI:** 10.64898/2026.10.02.756280

**Authors:** Alexis Hayes, Santiago Moreno-Caceres, Amber Delmas Eliason, David P. Marciano

## Abstract

Redox homeostasis requires both inducible antioxidant defense and constitutive machinery that continuously regenerates cellular reducing capacity. How these systems are altered across lung cell populations in chronic obstructive pulmonary disease (COPD) remains unclear. We integrated donor-level single-nucleus lung transcriptomes with smoke-exposure models, independent cohorts, and large-scale pharmacologic perturbation data. Both redox programs were reduced in established COPD, but constitutive redox support showed the greater loss, concentrated in fibroblasts and lymphatic endothelium. In fibroblasts, PGC-1α target-program output showed the most negative effect among 515 regulator programs despite preserved *PPARGC1A* expression. Smoke exposure activated inducible antioxidant defense across experimental models, whereas this adaptive program was reduced in established COPD. Pharmacologic perturbations further separated the two arms of redox homeostasis, with redox-directed interventions producing larger transcriptional responses in inducible defense and a distinct set of compounds increasing constitutive support together with PGC-1α target output and movement away from an independently defined COPD fibroblast state. These findings identify a cell-selective loss of constitutive redox support in COPD and implicate reduced PGC-1α target-program output in the fibroblast redox phenotype.

## Introduction

Oxidative stress in COPD is commonly viewed through the balance between oxidant burden and inducible antioxidant defense. Cellular redox homeostasis also depends on constitutive metabolic and enzymatic systems that sustain reducing capacity by generating NADPH and maintaining the glutathione, thioredoxin, and peroxiredoxin cycles, along with mitochondrial redox support. Whether this constitutive machinery is disrupted in COPD, and whether it behaves differently from inducible antioxidant defense, remains poorly resolved. COPD is a progressive, airflow-limiting disorder driven by chronic inhalational exposure [1], and oxidized lipids, nucleic acids, and proteins are detectable in diseased lung and circulation [2,3].

NRF2 is the canonical regulator of inducible antioxidant defense [4], which is rapidly engaged by oxidant exposure. We defined an inducible-defense program empirically from genes reproducibly activated by cigarette smoke in two independent human airway systems. This program captures the broader adaptive response to smoke and is analyzed separately from curated NRF2 regulon activity. Constitutive redox support represents a complementary arm of redox homeostasis, comprising the metabolic and enzymatic machinery that continuously regenerates NADPH and maintains glutathione, thioredoxin, peroxiredoxin, and mitochondrial redox systems. PGC-1α provides a plausible regulatory connection to this constitutive machinery through its coordinated control of mitochondrial metabolism and antioxidant support. Although the extent of NRF2 dysfunction varies across pulmonary compartments and studies, reduced NRF2 protein or NRF2-dependent antioxidant capacity has been reported in emphysematous lung and in pulmonary macrophages from patients with COPD and aged smokers [5,6,7]. Much of COPD redox biology and therapeutic development has focused on antioxidant supplementation and pharmacologic activation of inducible defense, leaving the machinery that sustains reducing capacity less well resolved [8,9,10].

Chronic redox dysfunction may also be highly specific to particular cell populations. Fibroblasts, endothelial cells, epithelial cells, and immune populations operate in different metabolic environments, and a defect restricted to one population can disappear in whole-lung averages [11]. PGC-1α transcriptional output can also change independently of *PPARGC1A* transcript abundance through localization, interacting partners, and post-translational regulation [12], so its target genes can report a regulatory deficit that transcript abundance does not.

We asked whether inducible antioxidant defense and constitutive redox support remain coupled in COPD when resolved across lung cell populations. Integrating donor-level single-nucleus transcriptomes with exposure models, independent cohorts, and orthogonal perturbation datasets identified a disproportionate loss of constitutive support in fibroblasts and lymphatic endothelium and a marked reduction of PGC-1α target-program output in COPD fibroblasts despite preserved *PPARGC1A* expression.

## Results

### Established COPD preferentially reduces constitutive redox support

Across the 12 lung populations meeting the analysis threshold (eight compartments and four subtypes, with fibroblasts and lymphatic endothelium represented at both levels), both redox programs shifted downward in COPD, but constitutive redox support declined more strongly in the populations showing the largest redox changes. The inducible-defense program was negative in all 12 populations and significant in five, so the preferential constitutive loss reflects a deeper deficit in constitutive support rather than preservation of inducible defense. The effect was strongest in fibroblasts and lymphatic endothelium, where the constitutive-minus-inducible contrasts were -0.26 SD in fibroblasts (95% CI -0.49 to -0.08, FDR = 0.044) and -0.22 SD in lymphatic endothelium (95% CI -0.34 to -0.09, FDR = 0.016, Figure 1A, 1B, Supplementary Table S2a). The contrast was negative in 11 of the 12 populations tested, so the pattern was not confined to the two populations reaching significance.

**Figure 1.**
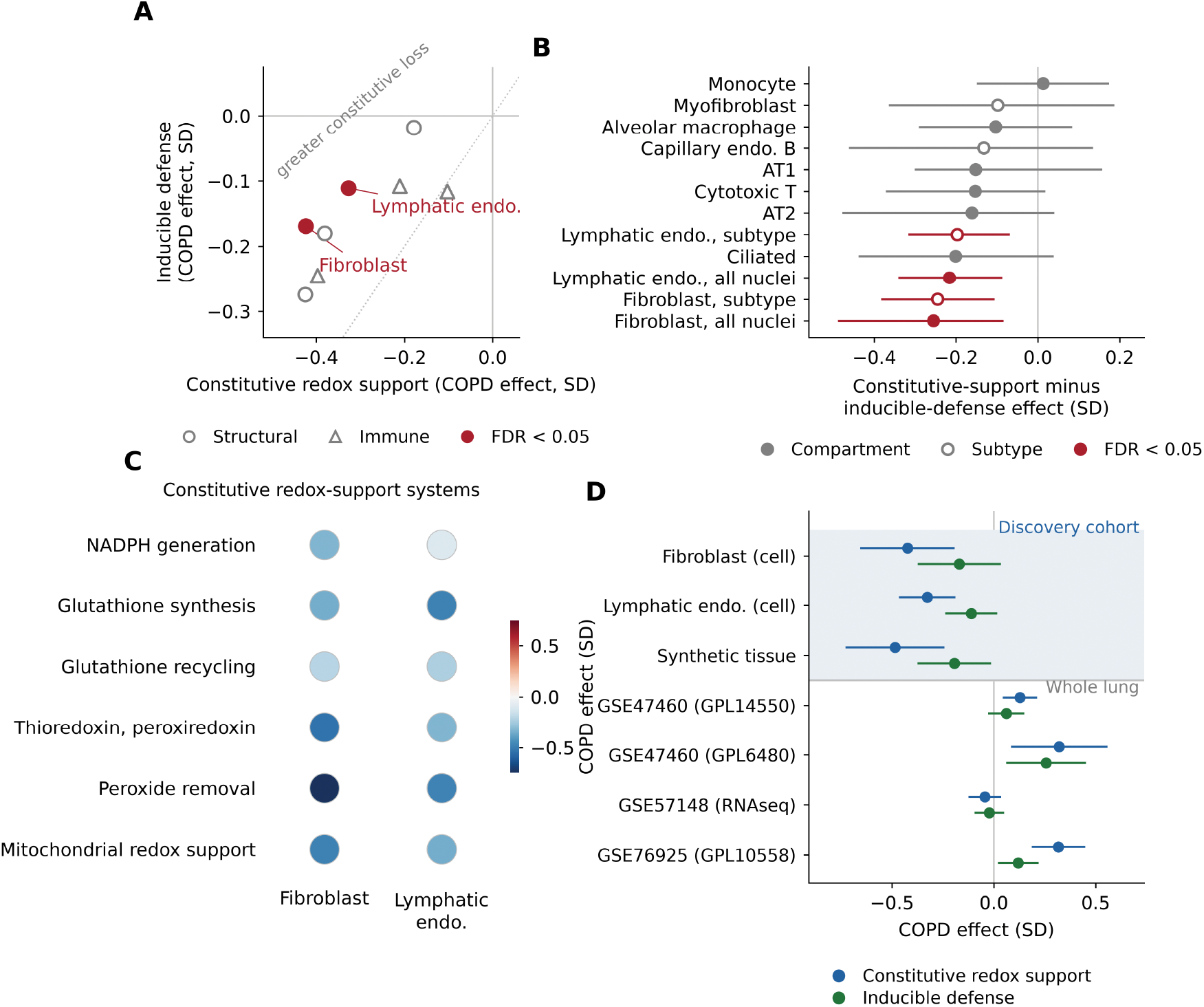
Established COPD preferentially reduces constitutive redox support. Our donor-level reanalysis of GSE136831 scored prespecified constitutive redox-support and inducible-defense programs across lung populations. (A) COPD effects on the two programs, both declining in the most affected structural populations while fibroblasts and lymphatic endothelium show the greater loss of constitutive support. (B) Constitutive-minus-inducible contrasts with 95% confidence intervals at compartment and subtype resolution, where negative values indicate a larger COPD-associated decline in constitutive redox support than in the inducible-defense program. (C) COPD effects across the six biochemical systems underlying constitutive support, in the two populations meeting the false-discovery criterion on the direct contrast, with color showing the signed donor-level effect and matched-null false-discovery values in Supplementary Table S2c. (D) The same two programs in the cell-resolved discovery populations, synthetic whole tissue reconstructed from the same nuclei, and three independent whole-lung cohorts. FDR values are Benjamini-Hochberg across tested populations. Datasets are GSE136831, GSE47460, GSE57148 and GSE76925. Endo., endothelial.

The constitutive program spans NADPH generation, glutathione synthesis and recycling, thioredoxin and peroxiredoxin cycling, peroxide removal and mitochondrial redox support, whereas the inducible program comprises genes reproducibly activated by cigarette smoke in human airway epithelium. The two programs share no genes; their construction is given in Methods, and the preferential constitutive deficit was preserved under two alternative definitions of the constitutive axis (Supplementary Figure S1, Tables S2a and S4).

The deficit was not confined to any single antioxidant pathway within the constitutive program. All six biochemical systems within the constitutive program shifted downward together in the two affected populations (Figure 1C, Supplementary Table S2c).

The deficit persisted when donor profiles were recombined into synthetic whole tissue (constitutive -0.49 SD, 95% CI -0.75 to -0.22, against -0.19 SD for inducible defense) but was not recovered in three independent whole-lung cohorts, where constitutive support was unchanged or higher in COPD (Figure 1D, Supplementary Tables S6a and S6b). Standardizing cellular composition within the discovery cohort preserved the effect, so the signal does not follow from differing cell proportions in that dataset (Supplementary Figure S1, Table S6b). Tissue averaging within the discovery cohort therefore does not explain the difference from the whole-lung cohorts, which remains unresolved.

Independent single-cell replication was limited by cell recovery and donor depth. Neither available cohort provided sufficient donor-level representation of both fibroblasts and lymphatic endothelium to retest the two strongest discovery signals. In the small emphysema cohort that recovered fibroblasts, the constitutive-minus-inducible contrast was concordant in direction at - 0.67 SD (95% CI -0.91 to -0.44) and negative across all eight annotated populations (Supplementary Figure S2, Tables S5a and S5b).

### COPD fibroblasts lose PGC-1α target-program output despite preserved *PPARGC1A* expression

The fibroblast constitutive-support deficit was accompanied by a marked reduction in PGC-1α target-program output. Across 515 regulator programs tested without preselection, PGC-1α showed the most negative COPD-associated effect and was the only regulator whose curated targets were also significantly enriched within the constitutive redox-support program (Figure 2A, Supplementary Tables S7a and S7b).

**Figure 2.**
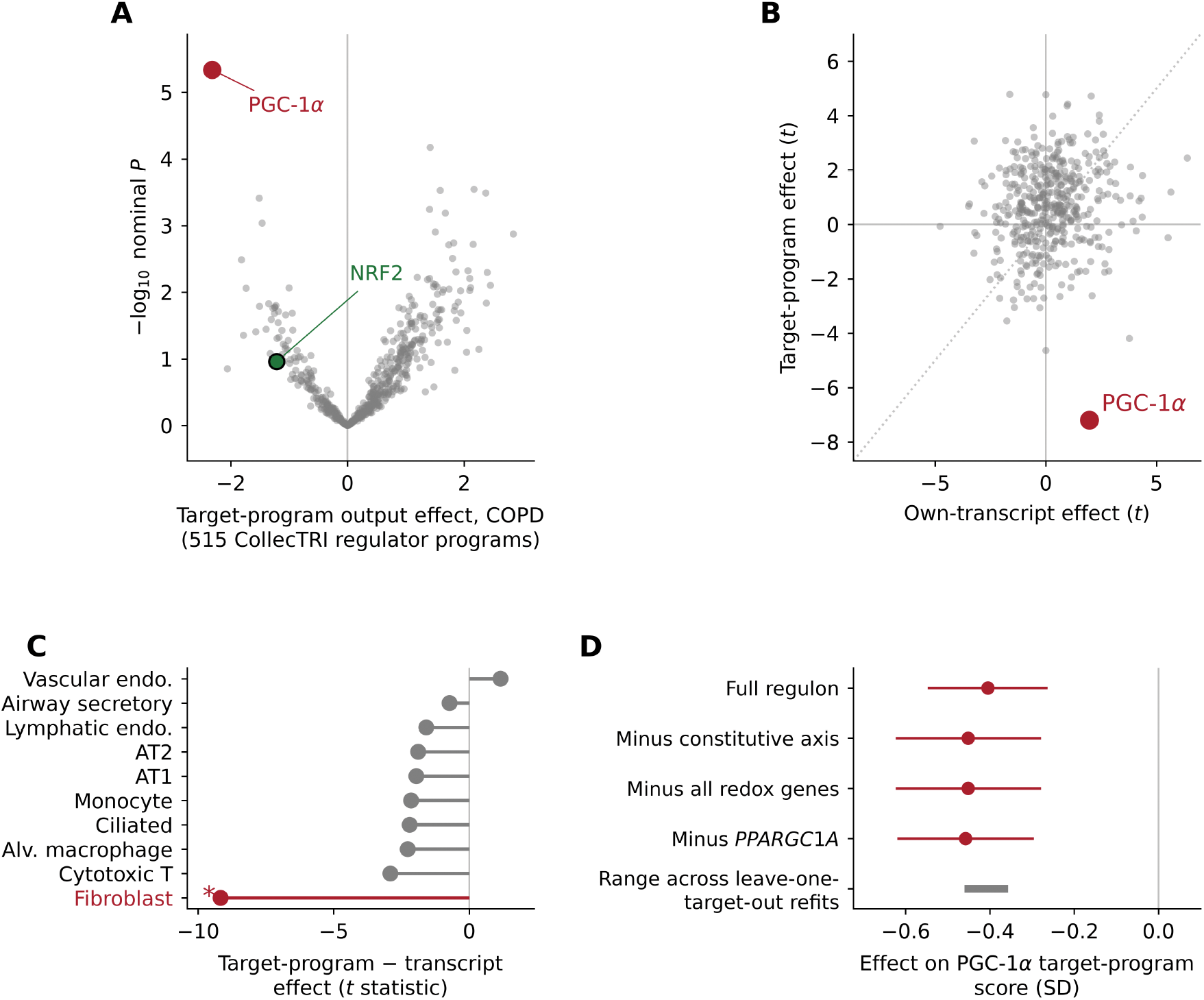
COPD fibroblasts lose PGC-1α target-program output despite preserved *PPARGC1A* expression. Our reanalysis of GSE136831 scored curated CollecTRI transcriptional programs at donor level. (A) COPD effects across 515 regulator programs. (B) Each regulator’s own-transcript effect against its target-program effect in the same cells and donors, with the dotted line marking agreement between the two. (C) Transcript-to-target discordance across lung populations, as the target-program effect minus the own-transcript effect. Fibroblasts are shown in red, and an asterisk denotes Benjamini-Hochberg FDR below 0.05 across tested populations. (D) PGC-1α target-program effect after removal of redox genes, of *PPARGC1A,* and of individual regulon targets, showing that the signal is robust to target-set composition. All three displayed confidence intervals exclude zero, and the gray bar spans the range across leave-one-target-out refits rather than a confidence interval. Activity denotes inferred transcriptional-program output rather than direct PGC-1α protein activation.

PGC-1α showed the most negative effect by both coefficient and *t* statistic, with an inferred-activity disease effect of -2.32 (SE 0.32, *t* = -7.19, *P* = 4.6 x 10^-6, FDR = 0.0024), and the next most negative programs, MYC, ZNF362, MXD1, VHL, and SREBF1, did not reach significance. Three of the 27 detected curated PGC-1α targets, *PPARGC1A*, *SOD2* and *TFAM*, fall within the constitutive redox-support axis (enrichment FDR = 0.0009).

*PPARGC1A* expression itself was preserved, separating coactivator transcript abundance from target-program output. Across 444 regulators measurable both ways in the same cells and donors and under one model, PGC-1α showed the largest transcript-to-target discordance, with a model *t* statistic of +1.97 for its own transcript (*P* = 0.068) against -7.19 for its target program (Figure 2B), and the effect was strongest in fibroblasts and reached significance only there (Figure 2C).

The decrease persisted after removal of the *PPARGC1A* autoregulatory edge, of constitutive-support and redox genes, and of every individual regulon target (Figure 2D, Supplementary Figure S3 and Table S7c). Two independent PGC-1 perturbation signatures were also concordant in direction with the COPD fibroblast state, since PGC-1α overexpression shifted fibroblast transcription away from that state while combined PGC-1α and PGC-1β depletion shifted it toward that state (Supplementary Table S7d). These results identify reduced PGC-1α target-program output as the strongest regulatory signature associated with the fibroblast redox-support deficit despite preserved *PPARGC1A* transcript abundance. The broader inducible-defense program was also lower in COPD fibroblasts, whereas curated NRF2 target-program activity was not significantly reduced in either regulon resource (*P* = 0.36 and 0.11), showing that the empirically defined smoke response and direct NRF2 regulon activity are related but distinct quantities.

### The smoke-induced defense program is reduced in established COPD

Cigarette smoke activated inducible antioxidant defense in two independent human airway systems, increasing NRF2 targets in both and glutathione metabolism in the air-liquid interface system [13,14]. The response was elevated at 7 days and again at 9 months of smoke exposure in the longitudinal mouse model and was lower after cessation than under continued exposure [15] (Figure 3A, 3B, Supplementary Table S3b). In established COPD, the same adaptive program was reduced in five of eight lung populations, including fibroblasts (*z* = -3.6, FDR = 0.003), cytotoxic T cells, AT1 cells, ciliated cells, and monocytes, while smoke-repressed genes tended to increase (Figure 3C). Across the ranked transcriptome, increasing smoke responsiveness was weakly associated with lower expression in several COPD lung populations, including fibroblasts (ρ = -0.118 across 9,334 genes, Figure 3D, Supplementary Tables S3c and S3d). A correlation of this size describes a shift in the ranked distribution rather than gene-for-gene reversal. In established COPD, the adaptive smoke-response program is reduced rather than maintained, alongside an even larger loss of constitutive redox support in fibroblasts and lymphatic endothelium.

**Figure 3.**
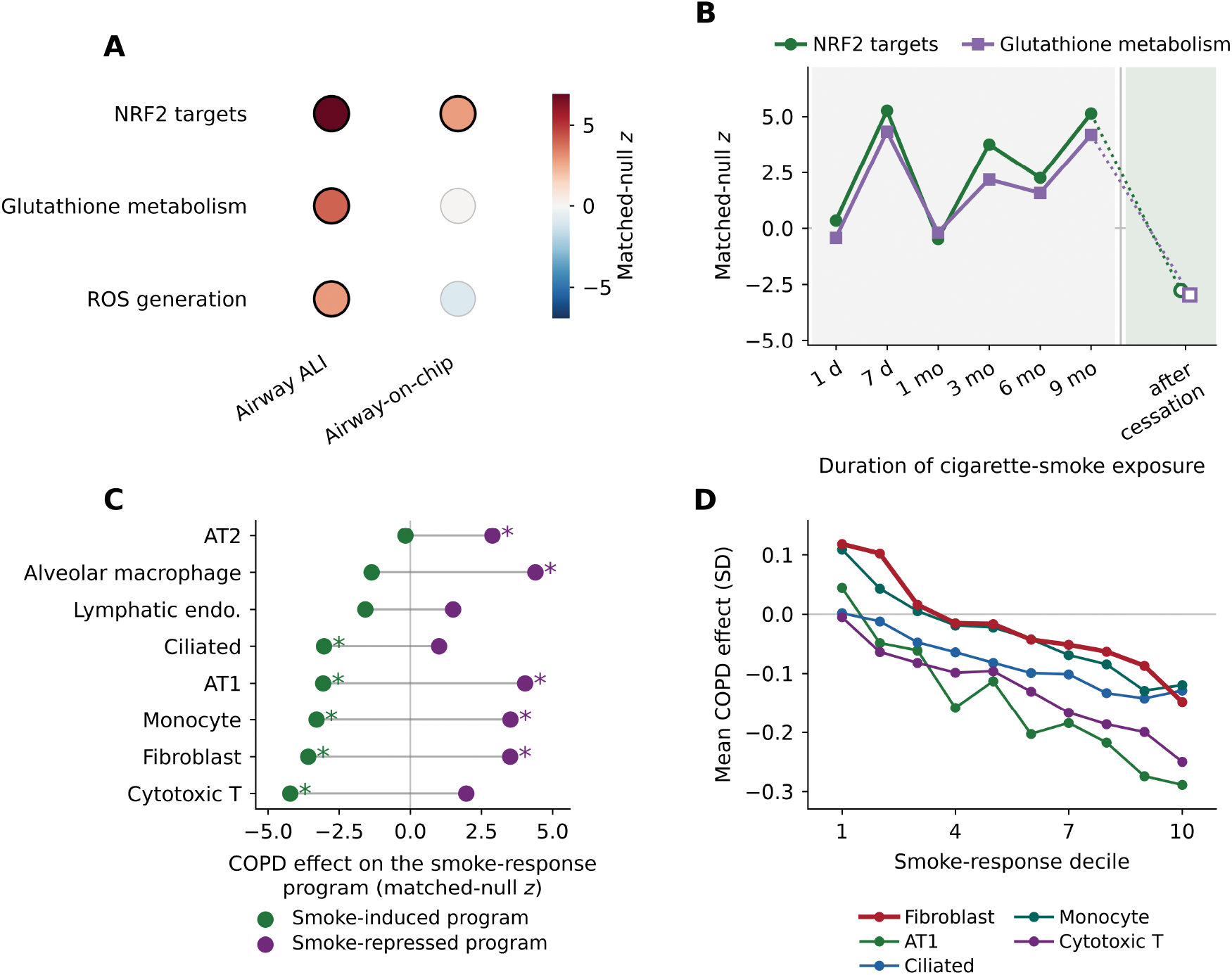
The smoke-induced defense program is reduced in established COPD. (A) We recalculated NRF2-target and glutathione responses to smoke from published datasets in two independent human airway systems, primary small-airway cells at air-liquid interface and bronchiolar epithelium in airway-on-chip devices. (B) The same programs during longitudinal mouse smoke exposure, each timepoint against air-exposed controls, and after cessation, shown as open symbols and contrasted against continued smoke exposure at 9 months [15]. (C) COPD effects on the genes reproducibly induced or repressed by smoke across lung populations, with the two estimates for one population joined by a gray line. An asterisk denotes program-level FDR below 0.05. (D) COPD effects across deciles of the ranked smoke response in the populations meeting the prespecified display criterion, five of eight tested, given with all eight in Supplementary Table S3d, with fibroblasts drawn heavier. The smoke-induced defense program is broader than a curated NRF2 regulon, and panels C and D report that program rather than direct NRF2 activity. In panel A, black outlines denote module-level FDR below 0.05. Datasets are GSE135188, GSE87098, GSE76205 and GSE136831.

### Redox-directed interventions produce larger responses in inducible defense

Across three direct redox perturbation datasets, inducible defense showed larger transcriptional responses than constitutive redox support, although the direction of the inducible response differed among interventions. Omaveloxolone, an NRF2-activating triterpenoid [16], produced a strong dose-dependent increase in inducible defense in primary human dermal fibroblasts, reaching +1.23 SD at matched-null *z* = 11.6 (FDR = 0.001) at 0.37 μM, while constitutive redox support did not differ from expression-matched null gene sets at any dose (competitive FDR ≥ 0.07) (Figure 4A, Supplementary Table S9a). A sulforaphane-class perturbation showed the same selectivity in an infected Calu-3 airway epithelial system [17] (Supplementary Table S9b). In a randomized double-blind placebo-controlled trial of nicotinamide riboside in patients with COPD [18], the inducible program shifted relative to expression-matched null gene sets (competitive *z* = -5.42, *P* = 0.0005) while the constitutive program did not (competitive *z* = -0.81, *P* = 0.42, Figure 4B, Supplementary Figure S5 and Table S9c).

**Figure 4.**
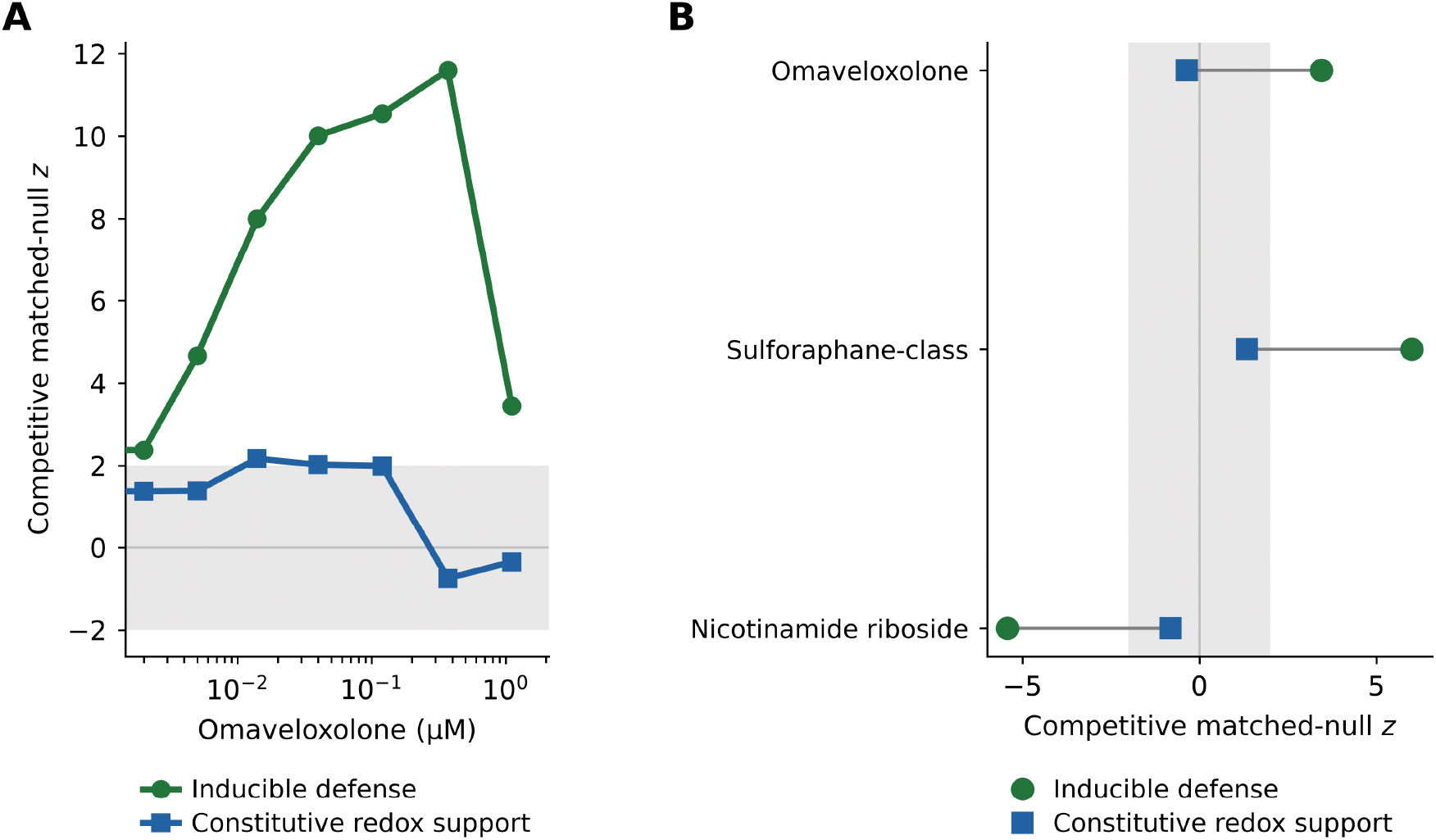
Redox-directed interventions produce larger responses in inducible defense than in constitutive redox support. Program scores are our reanalysis of published experiments. (A) Omaveloxolone dose response in primary human dermal fibroblasts, as competitive matched-null *z* for both programs, with shading marking the matched-null band. (B) Paired program effects across omaveloxolone-treated primary human dermal fibroblasts (highest dose passing read-depth QC, 1.11 μM), a sulforaphane-class perturbation in infected Calu-3 airway epithelium and nasal-brush transcriptomes from the randomized nicotinamide-riboside COPD trial, which was a patient trial rather than a cell system. The inducible program responds in different directions across these interventions, upward in the two cell systems and downward in the trial, while constitutive support remains comparatively stable. Gray shading marks a competitive matched-null *z* of magnitude below 1.96 in both panels. Datasets are GSE306317 and GSE243268, with the trial data at doi 10.5061/dryad.d2547d8b5.

The inducible response rose in the two cell systems and fell in the trial, so the common feature across these experiments was the greater responsiveness of the inducible program rather than any shared direction. The constitutive-minus-inducible contrast in the fibroblast dose series was negative and significant under all three axis definitions (*P* ≤ 0.0007). A strong inducible antioxidant response does not imply engagement of constitutive redox support.

### Chemical perturbation separates inducible defense from constitutive redox support

Across 107,404 chemical perturbation signatures representing 1,768 compounds, effects on constitutive redox support and inducible defense were weakly correlated (Pearson *r* = 0.23, *r*² = 0.05, Spearman ρ = 0.35), and 197 compounds significantly increased the constitutive program (Figure 5A, Supplementary Table S10d). The programs were separable in both directions, since 150 compounds raised constitutive support without raising inducible defense, 172 raised inducible defense alone, and 58 moved the two programs in opposite significant directions.

**Figure 5.**
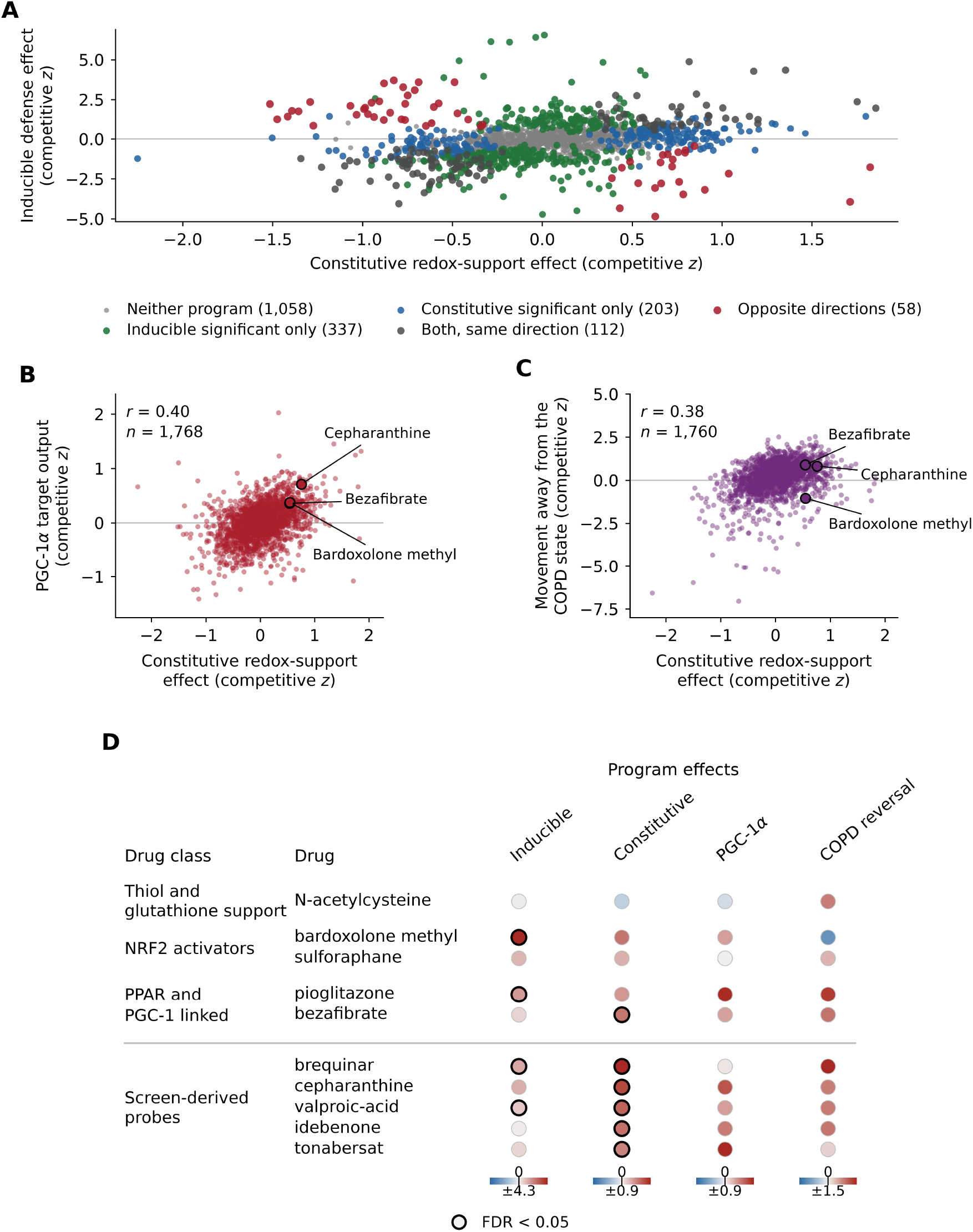
Chemical perturbation separates inducible defense from constitutive redox support. (A) Our reanalysis of 1,768 LINCS compounds, colored by which program moved significantly and in which direction. These categories include significant increases and decreases, whereas the positive-only counts in the Results refer to compounds that increased the program named. (B) Constitutive-support effect against PGC-1α target-program output. (C) The same effect against reversal of a gene-disjoint COPD fibroblast signature, where positive values denote movement away from the COPD state. The same three compounds are labeled in both. (D) A curated subset: the five named comparators discussed in the text above the rule and the five prespecified screen-derived probes below it, against the four program measures. Color is scaled independently within each column and centered at zero, with its range given beneath, and black outlines mark FDR below 0.05 for the two primary programs. Scores are from our reanalysis. Panel D is a curated subset of the complete prior-evidence inventory, which covers all 28 interventions and is given in Supplementary Figure S6 and Table S11.

Perturbations that increased constitutive redox support also tended to increase PGC-1α target output and to move fibroblast transcription away from the COPD state (*r* = 0.40, 95% CI 0.36 to 0.44, Figure 5B, and *r* = 0.38, 95% CI 0.34 to 0.42, Figure 5C). We constructed the COPD reversal signature to share no genes with the constitutive redox-support axis, so the second relationship is not a restatement of the first and indicates that an increase in the constitutive program accompanies broader transcriptional movement away from the disease state. Both relationships were essentially unchanged after controlling for cell-cycle score, transcriptional activity, and signature count (partial *r* = 0.37 and 0.37, Supplementary Table S10c).

To place this chemical landscape in the context of COPD pharmacology, we mapped redox-directed interventions identified from a prespecified literature and clinical-trial search onto the same transcriptional measures whenever compatible LINCS signatures were available (Figure 5D, Supplementary Figure S6 and Table S11). Interventions without compatible signatures remained in the evidence inventory but were not assigned perturbational effects, which limits direct comparison with much of the existing clinical literature. Among the named comparators, the strongest NRF2-directed response was concentrated in inducible defense, with bardoxolone methyl providing the clearest example. Bardoxolone methyl produced the strongest inducible-defense response among the named comparators (+4.32, FDR = 0.006) without significantly increasing constitutive support (+0.55, FDR = 0.22), illustrating that strong inducible antioxidant activation does not by itself raise the constitutive redox-support program.

The two PPAR-directed compounds in the comparator set, pioglitazone and bezafibrate, both lay in the region of perturbation space characterized by increased constitutive support and PGC-1α target output, although only bezafibrate reached significance on constitutive support. This pattern has precedent in fibrotic lung disease, where PGC-1α is stably repressed in primary IPF fibroblasts and rosiglitazone raised *PPARGC1A* expression and attenuated fibroblast activation [19]. Bezafibrate provides a mechanistically anchored PPAR and PGC-1 comparator, having been reported to raise PPAR and PGC-1α-associated mitochondrial biogenesis in differentiating human neural cells [20], and it increased constitutive support (+0.54, FDR = 0.040) together with PGC-1α target output (+0.36) and COPD fibroblast-state reversal (+0.89). The unbiased screen identified 115 compounds meeting the prespecified probe criteria, from which five mechanistically diverse examples were selected for display on the basis of their biochemical-arm dependence rather than effect magnitude (Figure 5D, Supplementary Table S12). Cepharanthine represents an NADPH-arm-dependent response, its constitutive effect falling from +0.79 on the full axis to - 0.03 when those genes are withheld, whereas tonabersat retains its effect when individual arms are removed, and bezafibrate is likewise largely unaffected (Supplementary Figure S4, Tables S10a, S10b and S12). These compounds provide mechanistically distinct tools to test whether increasing constitutive redox support can reverse the COPD fibroblast transcriptional redox phenotype.

## Discussion

COPD shows concurrent impairment of inducible antioxidant defense and the constitutive machinery that sustains reducing capacity, with the larger deficit concentrated in fibroblasts and lymphatic endothelium. The larger constitutive deficit spans all six prespecified systems underlying constitutive redox support, namely NADPH generation, glutathione synthesis and recycling, thioredoxin and peroxiredoxin cycling, peroxide removal and mitochondrial redox support, and it persisted in synthetic whole tissue from the discovery cohort but was not recovered in independent whole-lung cohorts, a discrepancy that tissue averaging does not explain and that requires replication at cell resolution.

In fibroblasts, the constitutive redox-support deficit is accompanied by reduced PGC-1α target-program output despite preserved *PPARGC1A* expression. This transcript-to-target discordance is consistent with regulation of PGC-1α output beyond coactivator transcript abundance, through localization, interacting partners, and post-translational modification [12], including NAD+-dependent deacetylation by SIRT1 [21]. SIRT1 protein has also been reported to be reduced in peripheral lung tissue from smokers and patients with COPD [22], raising altered SIRT1-dependent regulation as a possible contributor to reduced PGC-1α output in disease. We did not measure SIRT1 abundance or PGC-1α acetylation in the fibroblasts analyzed here, and the reported reduction was in lung tissue rather than fibroblasts, so this mechanism remains to be tested directly. The coactivator couples mitochondrial biogenesis to antioxidant and metabolic support, offering a plausible link between the transcriptional deficit and the bioenergetic and redox state of chronically activated fibroblasts [23]. The same coactivator is stably repressed in primary fibroblasts from patients with idiopathic pulmonary fibrosis, where restoring its expression attenuated fibroblast activation [19], placing the present observation within an established precedent in chronic lung disease.

The smoke-response analysis adds complementary evidence to this model. Acute and chronic smoke exposure activates inducible antioxidant defense, and in the longitudinal mouse model that response receded after cessation [15]. The cross-sectional COPD data are a separate disease state rather than the endpoint of that experimental time course, and in established COPD the inducible-defense genes are reduced rather than persistently elevated. That pattern is compatible with impaired chronic adaptation to oxidative stress, and with reports of reduced NRF2 protein and NRF2-dependent antioxidant capacity in COPD lung and pulmonary macrophages [5,6]. The empirically defined smoke-response program is broader than NRF2 itself, and direct NRF2 regulon activity was not significantly reduced in the fibroblast analysis, so the data indicate attenuation of the adaptive smoke-responsive program without requiring a model of uniformly reduced NRF2 activity.

Pharmacologic perturbation reinforces the distinction between inducible antioxidant defense and constitutive redox support. Redox-directed interventions produce larger transcriptional responses in the inducible program, whereas the large chemical screen identifies a different set of perturbations that increase constitutive support. Those effects track with both PGC-1α target-program output and movement away from an independent COPD fibroblast signature, and because that signature shares no genes with the constitutive redox-support axis, the relationship places constitutive redox support within a broader disease-associated transcriptional state rather than an isolated gene-set score. Inducible defense and constitutive redox support represent complementary pharmacodynamic targets rather than interchangeable measures of antioxidant activity.

The critical next step is to determine whether increasing constitutive redox support can restore functional redox capacity in primary COPD fibroblasts. The mechanistically distinct perturbations identified here can be tested alongside an inducible-defense activator while measuring PGC-1α target output, NADPH/NADP+, GSH/GSSG, and oxidative burden to test whether the constitutive redox-support deficit is functionally important and whether it can be corrected independently of inducible-defense activation. Direct measurement of SIRT1 abundance or activity and of PGC-1α acetylation could determine whether that regulatory route contributes to the fibroblast phenotype.

## 4. Limitations

Independent replication of the cell-resolved phenotype remains incomplete in the available cohorts. Neither available independent single-cell cohort recovered fibroblasts and lymphatic endothelium at the donor depth needed to test the two strongest discovery populations, and the one fibroblast-containing cohort was a small emphysema series with three donors per group. That comparison is concordant in direction rather than a definitive replication, and it carries both severity and a probable smoking imbalance.

The smoke-exposure experiments and the COPD lung transcriptomes were measured in different experimental systems, so the comparison does not establish a temporal trajectory from smoking to established disease; it contrasts states rather than a time course. The discovery metadata also lack smoking status, pack-years, and spirometric fields, so no model adjusted for smoking exposure could be fitted. A COPD-associated signal in these data cannot be separated from the cumulative smoking exposure that accompanies the disease, and the smoke-exposure experiments address the acute response in airway epithelium rather than the chronic state of fibroblasts.

Because the two redox programs are derived from gene expression rather than direct measurements of NADPH, glutathione, or thiol status, the present study defines a transcriptional redox state rather than biochemical reducing capacity. Regulator activity is likewise inferred from curated signed target expression rather than measured as protein abundance, chromatin occupancy, or post-translational state, and we measured neither SIRT1 abundance nor PGC-1α acetylation.

The chemical-perturbation analysis is also constrained by available signatures, because many redox-directed agents tested clinically in COPD are absent from the library and most available compound signatures derive from non-COPD cell systems. Only one of the named comparators carries enough signatures from a fibroblast, stromal, lung, airway, or endothelial context to support a context-restricted analysis (Supplementary Table S11). The screen identifies transcriptional behavior and experimental probes rather than therapeutic efficacy, and this does not imply that antioxidant or mucolytic therapies lack clinical benefit [8,9], since those agents can act through extracellular thiol chemistry, mucus handling, or inflammatory pathways.

## Methods

### Discovery single-cell data and pseudobulk

We used single-nucleus profiles from 46 donors from the lung cell atlas GSE136831, generated and deposited by Adams and colleagues [24], and including control, COPD, and idiopathic pulmonary fibrosis donors. We used only the 18 COPD and 28 control donors, and Sauler and colleagues describe their COPD phenotype in the disease-specific analysis of the same resource [11]. Raw UMI counts were summed per donor and cell type, and all statistics were computed at donor level. Compartments required at least 20 nuclei per donor and 6 donors per group, and subtypes at least 10 nuclei per donor and 8 donors per group, leaving 77,823 nuclei in 264 groups from 45 donors. We converted counts to log_2_ CPM, standardized per gene across donors, and included age, sex, and log_10_ pseudobulk cell number in every model.

### Module testing

Module shifts were assessed both self-contained and against 2,000 expression-matched null gene sets of equal size drawn from decile-matched expression bins. Disease coefficients provide the effect estimates in the text and figures, and the null sets test whether a program moves beyond genes of comparable abundance, supplying the competitive statistics reported as matched-null *z*.

### Axis definitions

Redox programs were defined before any disease analysis and then held fixed. The inducible-defense axis comprises the 200 genes most reproducibly increased by whole cigarette smoke, requiring concordant direction in both independent human airway exposure systems, so it captures the acute smoke-responsive transcriptional program empirically rather than by annotation. The primary constitutive redox-support axis was built in three declared steps. It began with an a priori collection of genes encoding machinery for NADPH generation, glutathione synthesis and recycling, thioredoxin and peroxiredoxin cycling, peroxide removal, and mitochondrial redox support, from which every gene induced concordantly in both smoke systems was removed, leaving 41 genes with no overlap with the inducible axis (Supplementary Table S1). Membership was not modified for any later dataset. Module-level results are reported on the six prespecified parent modules rather than on the filtered axis. Because removing smoke-responsive genes could in principle manufacture the distinction, all primary analyses were repeated with the complete a priori set and with a score residualized against the inducible axis.

### Direct axis contrast

For each population, the per-donor difference between the constitutive and inducible axis scores was regressed on disease status with the same covariates. The disease coefficient equals the difference between the two separately fitted axis coefficients, and its standard error accounts for covariance between the scores. Uncertainty is reported as a model standard error, a donor-bootstrap 95% CI and a label-permutation p-value, with Benjamini-Hochberg FDR across populations.

### Axis-definition sensitivity

Two alternative constitutive-axis definitions that do not use smoke response for gene selection were run through the same models: the full a priori set with no genes removed and that set residualized on the inducible score across donors. The primary axis was not redefined.

### Whole-ranked analysis

Gene-level COPD coefficients were obtained for all genes simultaneously by least squares on the shared design matrix. The smoke-response score is the mean of the two airway systems’ log-fold changes after standardizing each. The relationship was assessed by Spearman correlation across all shared genes and by decile trend, and the reproducibly smoke-induced and smoke-repressed programs were projected into each population against expression-matched nulls (Table S3a).

### Independent cohort

GSE279570 comprises 11 patients with mild to moderate COPD and 7 age-matched controls profiled by single-cell RNA-seq, with all rules fixed before analysis. Quality filtering, clustering, and marker-based label assignment followed the parameters given in Supplementary Methods. Populations were tested only with at least 20 cells per donor and 4 donors per group. Age and sex are not deposited for this series, so models contain disease status and log_10_ cell number, and populations below threshold are reported as untested rather than estimated at a relaxed threshold.

### Exposure datasets

GSE135188 provided primary human small-airway epithelium at air-liquid interface from 6 donors with whole smoke against air, paired within donor. GSE87098 provided a small-airway-on-chip model of healthy and COPD epithelium in a two-by-two design. GSE76205 provided mouse lung from one day to nine months of smoke exposure with a cessation arm, analyzed at module level only.

### Bulk cohorts and synthetic tissue

We analyzed GSE47460 separately by array platform, alongside GSE76925 [25] and GSE57148 [26], using available covariates, and reported each stratum rather than pooling. We recombined donor cell-type profiles into synthetic whole tissue at each donor’s observed composition and at a pooled-control reference composition to test whether cellular composition within the discovery resource accounts for the cell-resolved effect.

### Program terminology

The PGC-1α target program scored in the fibroblast regulator analysis and the PGC-1α measure scored in the compound screen are the same 27 curated genes, verified by direct set comparison, so the two results reflect one program rather than two similar ones. Three further quantities are reported separately and are not interchangeable. The inducible-defense program is the empirically defined set of genes reproducibly activated by cigarette smoke in the two human airway systems, the NRF2 regulon is the curated NFE2L2 target program scored from regulon resources, and inducible antioxidant defense is the broader biological concept that the inducible-defense program approximates. The inducible-defense program is NRF2-centered but is not an NRF2 regulon, and the two are scored independently throughout.

### Regulator activity inference

Regulator activity was inferred per donor from curated signed target sets using a univariate linear model over two regulon resources, DoRothEA confidence levels A to C [27] and CollecTRI [28], with decoupleR [29] and a minimum of ten detected targets per regulon. Donor-standardized expression was the input, so activity reflects relative donor differences rather than expression scale. The 515-program discovery screen is the CollecTRI screen, since that resource contains a PGC-1α regulon, and DoRothEA scored 268 programs as an additional resource covering *NFE2L2*. No candidate regulator was preselected. The COPD effect on each regulon was fitted at donor level with age, sex, and log_10_ pseudobulk cell number as covariates, matching every other model reported here, with Benjamini-Hochberg FDR [30] applied within each cell type and resource. The dual criterion required both reduced inferred activity and hypergeometric enrichment of curated targets among the constitutive redox-support axis, against the detected transcriptome as background, and we repeated both analyses with the PGC-1α autoregulatory edge removed. We assessed circularity by refitting activity after removing the constitutive-axis genes from the regulon, and again after removing every redox-module gene. Two estimators are reported and are not on a common scale: the primary decoupleR score over the signed regulon and an independently recomputed signed target score in expression standard deviations that allows individual targets to be removed deterministically. For every regulator with both a detected transcript and a scored regulon in the same population, we estimated transcript and inferred-activity effects from the same model, and defined discordance as the activity statistic minus the transcript statistic, then placed it in the empirical distribution across all regulators measurable both ways. Estimator variants and leave-one-target-out refitting are described in Supplementary Methods.

### Redox-directed perturbations

GSE306317 provided primary human dermal fibroblasts treated with omaveloxolone across ten log-spaced doses against 32 vehicle wells. We retained wells with more than one million assigned reads, excluding the two highest doses, and verified target engagement beforehand with a canonical NRF2 marker panel that contributed to neither axis. GSE243268 provided a stabilized sulforaphane preparation against untreated control in infected Calu-3 airway epithelial cells at two time points; we analyzed it within time point and treated it as supporting rather than primary evidence because the system is neither a fibroblast nor an uninfected epithelium. Axis effects were tested self-contained and against 2,000 abundance-matched null gene sets; the between-axis contrast was estimated on the per-sample difference of the two scores, and both alternative constitutive-axis definitions were run as sensitivity analyses.

### Randomized nicotinamide riboside trial

Individual-level nasal brush RNA-seq from the ICON trial (NCT04990869, Dryad 10.5061/dryad.d2547d8b5) provided normalized counts for 143 samples from 39 participants with COPD and 17 controls at baseline, post-treatment, and follow-up. Counts were log-transformed, filtered to genes detected in over half of samples, and standardized per gene. We randomized every contrast, first computing within-subject change and then comparing arms with a Welch t-test; we never reported within-arm change alone. The competitive test permuted 2,000 abundance-matched gene sets through the identical contrast.

### Compound-library screen

Level 5 consensus signatures from GSE70138 were scored with the same rank-based directional statistic used throughout. Of 118,050 signatures, the 107,404 annotated as compound perturbations entered scoring, spanning 1,768 compounds, with axis definitions fixed beforehand and scores oriented so that positive denotes raising the genes COPD reduces. Null matching was stratified on L1000 landmark status, cross-signature standard deviation, and mean absolute z. Because the raw constitutive-support score retained an association with proliferation and global transcriptional activity, we residualized every axis on the cell-cycle module score, the transcriptional activity score, and signature strength, and classified every compound on the adjusted statistic alone. Residualization materially changed the constitutive-positive set, reducing it from 357 compounds before adjustment to 197 afterward, and both statistics are distributed in Source Data. Signatures were aggregated to compounds as the median per-signature z with a 2,000-replicate bootstrap interval, requiring at least five signatures per compound, with Benjamini-Hochberg correction across compounds within each axis. Two measures defined outside the constitutive redox-support axis tested the screen against the disease analysis: the externally curated 27-gene PGC-1α target set, which shares *PPARGC1A*, *SOD2,* and *TFAM* with the primary axis, and a COPD fibroblast signature of the 200 most raised and 200 most reduced genes from which every a priori constitutive redox-support gene was removed, making it gene-disjoint by construction. Both were scored with the same statistic and null construction as the axes. A leave-one-biochemical-arm-out analysis removed each of the six arms in turn and repeated the entire procedure, with the full axis rescored in the same pass so that baseline and variants share one set of gene strata. We selected screen-derived probes using criteria fixed before examining any compound name, requiring a significant positive constitutive-support effect on the adjusted statistic, concordant increases in PGC-1α target output and fibroblast-state reversal, at least ten contributing signatures, and a complete leave-one-arm record. Of 115 eligible compounds, five are displayed, chosen to span arm-dependence profiles rather than to maximize any effect, and the full set with every criterion evaluated is in Supplementary Table S12. Bezafibrate was included separately as a mechanistically anchored PPAR and PGC-1 comparator. Omaveloxolone entered the inventory because we analyzed a dedicated dose series here rather than through the literature search. No compound was ranked or nominated therapeutically. Pearson correlation confidence intervals were obtained by Fisher’s z transformation.

### Redox-intervention evidence inventory

The prespecified literature and clinical-trial search returned 25 interventions, to which omaveloxolone, bezafibrate, and cepharanthine were added on the separate grounds described in the preceding section, for a total of 28. Fourteen of the 28 have a compatible signature in the perturbation library and carry program scores, and we retain all 28 with their evidence in the complete inventory in Supplementary Figure S6 and Table S11. Figure 5D displays a curated subset: the five named comparators discussed in the Results together with five prespecified screen-derived probes selected to span biochemical-arm dependence. The published literature and the ClinicalTrials.gov registry were searched before any compound was scored, using prespecified queries recorded in Source Data. We retained interventions when we could identify an administered redox-directed agent in a COPD, emphysema, or cigarette-smoke context, grouped by broad redox mechanism, and matched it to the perturbation library by recognized name and synonyms. Agents with no compatible library signature were classified as not scoreable rather than as null, and each absence was checked against the library name index.

### Multiplicity and software

Benjamini-Hochberg FDR was applied within analysis families, and all random procedures used fixed seeds. Analyses used Python 3.11.15, NumPy 2.4.6, pandas 2.3.3, SciPy 1.17.1, statsmodels 0.14.6, and Matplotlib 3.11.0. Figure-generation scripts and the final gene-set definitions accompany the submission.

## Supplementary materials

Supplementary Methods, Supplementary Results, Supplementary Figures S1 to S6 and Supplementary Tables S1 to S12 accompany the manuscript. The supplementary tables are supplied as a single workbook with an index sheet.

## Author contributions

All authors performed analyses and edited the manuscript. D.P.M. wrote the manuscript.

## Ethics

This study analyzed only previously published, de-identified human datasets available under their original consent and approval. No new human subjects were recruited, and no new samples were collected, so no additional institutional review board approval or informed consent was required. The accession and provenance of every dataset are given in Supplementary Table S8.

## Data availability

All primary data analyzed in this study are public, and Supplementary Table S8 provides the complete dataset and sample inventory, including assay type, scope, and point of use. Discovery single-nucleus data, GSE136831. Independent single-cell cohort, GSE279570. Human airway smoke-exposure experiments, GSE135188 and GSE87098. Longitudinal mouse smoke and cessation data, GSE76205 [15]. Bulk lung cohorts, GSE47460, GSE76925 [25] and GSE57148 [26]. Redox-directed perturbations: GSE306317 and GSE243268 [17]. GSE306317, GSE279570, and GSE70138 carry no associated primary publication and are cited by accession. PGC-1 perturbation experiments: GSE51498 and GSE108674. Emphysema single-cell cohort, GSE196638 [31]. Chemical-perturbation transcriptomes, GSE70138. The randomized nicotinamide riboside trial data are deposited at Dryad under doi:10.5061/dryad.d2547d8b5. Row-level data underlying every data-bearing main figure and Supplementary Figure S2 are provided as Source Data, distributed as Source_Data.zip. Figures 1 and 4 are backed by Source_Data_Figure1_panel_values.csv and Source_Data_Figure4_panel_values.csv, Figures 2, 3, and 5 by their own named Source Data files, and every displayed value also traces to a Supplementary Table through the figure source index distributed with the reproducibility archive. For the discovery cohort GSE136831, the deposited donor characteristics include tissue, disease status, age, and sex, with no smoking, pack-year, or spirometric field in the sample records or the cell-level metadata table, so we report no smoking-adjusted sensitivity model.

## Code availability

Figure-generation scripts, the gene-set and axis definitions, the machine-readable derived results, and a pinned computational environment specification accompany the manuscript as supplementary files, and every figure in this paper can be regenerated from the supplied package alone. The analysis code that produced the derived result tables from the primary datasets is available from the corresponding author on request. Analyses were run in Python 3.11.15, NumPy 2.4.6, pandas 2.3.3, SciPy 1.17.1, statsmodels 0.14.6, and Matplotlib 3.11.0. The supplied environment file reproduces the distributed figures from the derived tables and pins the packages those scripts require. It is not the complete primary-analysis environment. The single-cell and regulon analyses additionally used scanpy and decoupleR, which the figure scripts do not import and which the supplied environment does not pin.

## References

1. Global Initiative for Chronic Obstructive Lung Disease. Global Strategy for the Diagnosis, Management and Prevention of Chronic Obstructive Pulmonary Disease: 2026 Report. Deer Park, IL: GOLD, 2025. Available from goldcopd.org

2. Rahman I, Adcock IM. Oxidative stress and redox regulation of lung inflammation in COPD. Eur Respir J 2006;28:219–242. doi:10.1183/09031936.06.00053805

3. Yang S, Wu H, Zhao J, et al. Feasibility of 8-OHdG formation and hOGG1 induction in PBMCs for assessing oxidative DNA damage in the lung of COPD patients. Respirology 2014;19:1183–1190. doi:10.1111/resp.12378

4. Suzuki T, Yamamoto M. Stress-sensing mechanisms and the physiological roles of the Keap1-Nrf2 system during cellular stress. J Biol Chem 2017;292:16817–16824. doi:10.1074/jbc.R117.800169

5. Goven D, Boutten A, Lecon-Malas V, et al. Altered Nrf2/Keap1-Bach1 equilibrium in pulmonary emphysema. Thorax 2008;63:916–924. doi:10.1136/thx.2007.091181

6. Suzuki M, Betsuyaku T, Ito Y, et al. Down-regulated NF-E2-related factor 2 in pulmonary macrophages of aged smokers and patients with chronic obstructive pulmonary disease. Am J Respir Cell Mol Biol 2008;39:673–682. doi:10.1165/rcmb.2007-0424oc

7. Harvey CJ, Thimmulappa RK, Sethi S, et al. Targeting Nrf2 signaling improves bacterial clearance by alveolar macrophages in patients with COPD and in a mouse model. Sci Transl Med 2011;3:78ra32. doi:10.1126/scitranslmed.3002042

8. Decramer M, Rutten-van Mölken M, Dekhuijzen PN, et al. Effects of N-acetylcysteine on outcomes in chronic obstructive pulmonary disease (Bronchitis Randomized on NAC Cost-Utility Study, BRONCUS): a randomised placebo-controlled trial. Lancet 2005;365:1552–1560. doi:10.1016/S0140-6736(05)66456-2

9. Zheng JP, Wen FQ, Bai CX, et al. Twice daily N-acetylcysteine 600 mg for exacerbations of chronic obstructive pulmonary disease (PANTHEON): a randomised, double-blind placebo-controlled trial. Lancet Respir Med 2014;2:187–194. doi:10.1016/S2213-2600(13)70286-8

10. Wise RA, Holbrook JT, Criner G, et al. Lack of effect of oral sulforaphane administration on Nrf2 expression in COPD: a randomized, double-blind, placebo controlled trial. PLoS One 2016;11:e0163716. doi:10.1371/journal.pone.0163716

11. Sauler M, McDonough JE, Adams TS, et al. Characterization of the COPD alveolar niche using single-cell RNA sequencing. Nat Commun 2022;13:494. doi:10.1038/s41467-022-28062-9

12. Puigserver P, Spiegelman BM. Peroxisome proliferator-activated receptor-gamma coactivator 1 alpha (PGC-1 alpha): transcriptional coactivator and metabolic regulator. Endocr Rev 2003;24:78–90. doi:10.1210/er.2002-0012

13. Gindele JA et al. Intermittent exposure to whole cigarette smoke alters the differentiation of primary small airway epithelial cells in the air-liquid interface culture. Sci Rep 2020;10:6257. doi:10.1038/s41598-020-63345-5

14. Benam KH et al. Matched-Comparative Modeling of Normal and Diseased Human Airway Responses Using a Microengineered Breathing Lung Chip. Cell Syst 2016;3:456–466.e4. doi:10.1016/j.cels.2016.10.003

15. Miller MA et al. Gene and metabolite time-course response to cigarette smoking in mouse lung and plasma. PLoS One 2017;12:e0178281. doi:10.1371/journal.pone.0178281

16. Lynch DR, Chin MP, Delatycki MB, et al. Safety and efficacy of omaveloxolone in Friedreich ataxia (MOXIe study). Ann Neurol 2021;89:212–225. doi:10.1002/ana.25934

17. Romeo PH et al. Multiple Mechanisms of Action of Sulfodyne(®), a Natural Antioxidant, against Pathogenic Effects of SARS-CoV-2 Infection. Antioxidants (Basel) 2024;13. doi:10.3390/antiox13091083

18. Norheim KL, Ben Ezra M, Heckenbach I, et al. Effect of nicotinamide riboside on airway inflammation in COPD: a randomized, placebo-controlled trial. Nat Aging 2024;4:1772–1781. doi:10.1038/s43587-024-00758-1

19. Caporarello N, Meridew JA, Jones DL, et al. PGC1α repression in IPF fibroblasts drives a pathologic metabolic, secretory and fibrogenic state. Thorax 2019;74:749–760. doi:10.1136/thoraxjnl-2019-213064

20. Augustyniak J, Lenart J, Gaj P, et al. Bezafibrate upregulates mitochondrial biogenesis and influence neural differentiation of human-induced pluripotent stem cells. Mol Neurobiol 2019;56:4346–4363. doi:10.1007/s12035-018-1368-2

21. Rodgers JT, Lerin C, Haas W, Gygi SP, Spiegelman BM, Puigserver P. Nutrient control of glucose homeostasis through a complex of PGC-1alpha and SIRT1. Nature 2005;434:113–118. doi:10.1038/nature03354

22. Rajendrasozhan S, Yang SR, Kinnula VL, Rahman I. SIRT1, an antiinflammatory and antiaging protein, is decreased in lungs of patients with chronic obstructive pulmonary disease. Am J Respir Crit Care Med 2008;177:861–870. doi:10.1164/rccm.200708-1269OC

23. St-Pierre J, Drori S, Uldry M, et al. Suppression of reactive oxygen species and neurodegeneration by the PGC-1 transcriptional coactivators. Cell 2006;127:397–408. doi:10.1016/j.cell.2006.09.024

24. Adams TS, Schupp JC, Poli S, et al. Single-cell RNA-seq reveals ectopic and aberrant lung-resident cell populations in idiopathic pulmonary fibrosis. Sci Adv 2020;6:eaba1983. doi:10.1126/sciadv.aba1983

25. Morrow JD et al. Functional interactors of three genome-wide association study genes are differentially expressed in severe chronic obstructive pulmonary disease lung tissue. Sci Rep 2017;7:44232. doi:10.1038/srep44232

26. Kim WJ, Lim JH, Lee JS, et al. Comprehensive analysis of transcriptome sequencing data in the lung tissues of COPD subjects. Int J Genomics 2015;2015:206937. doi:10.1155/2015/206937

27. Garcia-Alonso L, Holland CH, Ibrahim MM, et al. Benchmark and integration of resources for the estimation of human transcription factor activities. Genome Res 2019;29:1363–1375. doi:10.1101/gr.240663.118

28. Müller-Dott S, Tsirvouli E, Vazquez M, et al. Expanding the coverage of regulons from high-confidence prior knowledge for accurate estimation of transcription factor activities. Nucleic Acids Res 2023;51:10934–10949. doi:10.1093/nar/gkad841

29. Badia-i-Mompel P, Vélez Santiago J, Braunger J, et al. decoupleR: ensemble of computational methods to infer biological activities from omics data. Bioinform Adv 2022;2:vbac016. doi:10.1093/bioadv/vbac016

30. Benjamini Y, Hochberg Y. Controlling the false discovery rate: a practical and powerful approach to multiple testing. J R Stat Soc Series B 1995;57:289–300. doi:10.1111/j.2517-6161.1995.tb02031.x

31. Wang C et al. Dysregulated lung stroma drives emphysema exacerbation by potentiating resident lymphocytes to suppress an epithelial stem cell reservoir. Immunity 2023;56:576–591.e10. doi:10.1016/j.immuni.2023.01.032

