## Supplemental_Information for "COPD Preferentially Disrupts Constitutive Redox Support Relative to Inducible Antioxidant Defense"

### Supplementary Methods

**PGC-1α robustness analyses.** Robustness analyses tested alternative signed target-score definitions, removal of the *PPARGC1A* autoregulatory edge, removal of constitutive-support or redox genes, and leave-one-target refits. Signed estimates remained negative across all of these analyses. Complete target-level sensitivity results are provided in Source Data.

**Independent cohort processing.** Cells in GSE279570 were retained with at least 500 detected genes and under 20% mitochondrial counts. Clustering used 2,000 highly variable genes, 50 principal components, and Leiden clustering at resolution 1.0; a cluster received a discovery label only when the best-matching canonical marker panel exceeded a fixed score margin. Age and sex are not deposited for this series, so models contain disease status and log10 cell number, and populations below threshold are reported as untested rather than estimated at a relaxed threshold.

**Leave-one-biochemical-arm analysis.** Each of the six biochemical parent modules was removed in turn from the constitutive-support axis, every compound was rescored, and the change relative to the full axis was used to identify biochemical-arm dependence. We rescored the full axis within the same matched analysis set, producing minor differences from the values in Figure 5.

**Redox-intervention evidence inventory.** We searched the published literature and the ClinicalTrials.gov registry before scoring any compound against the perturbation library. Retained interventions were those for which an administered redox-directed agent could be identified in a COPD, emphysema, or cigarette-smoke context. Interventions were grouped by broad redox mechanism and matched to the perturbation library using recognized compound names and synonyms. Compounds without a compatible signature were classified as not scoreable rather than as null effects. The complete search record and the alias-resolution audit are provided in the reproducibility archive.

### Supplementary Results

**Axis-definition sensitivity.** The complete a priori constitutive set, with no genes removed for smoke responsiveness, reproduced the preferential loss of constitutive-support transcription in COPD. A constitutive score residualized against the inducible axis reproduced the same pattern. Fibroblasts and lymphatic endothelium remained among the most strongly affected populations under both definitions (Supplementary Figure S1, Table S4).

**Whole-tissue composition.** Recombining the same donor cell profiles into synthetic whole tissue, at each donor's observed cellular composition and at a standardized reference composition, retained the larger loss of constitutive support at the whole-tissue level (Supplementary Figure S1, Table S6b).

**Independent cohort support.** The independent COPD single-cell cohort did not recover fibroblasts or lymphatic endothelium at the donor depth needed to test the two strongest discovery populations. In the small emphysema cohort that did contain fibroblasts, the difference between the two programs was concordant in direction with the discovery result. With three donors per group, this is supportive rather than independent replication (Supplementary Figure S2, Tables S5a and S5b).

**Independent PGC-1 perturbation support.** PGC-1α overexpression produced a signature running counter to the COPD fibroblast ranking, while combined PGC-1α and PGC-1β depletion produced one resembling it. Neither perturbation was performed in lung fibroblasts, so these observations are directional support rather than replication (Supplementary Figure S3, Table S7d).

### Supplementary Figures

**Abbreviations used in the supplementary figures.** Endo., endothelial. Alv., alveolar. NK, natural killer. GSH, glutathione. TXN, thioredoxin. PRDX, peroxiredoxin. ALI, air-liquid interface.

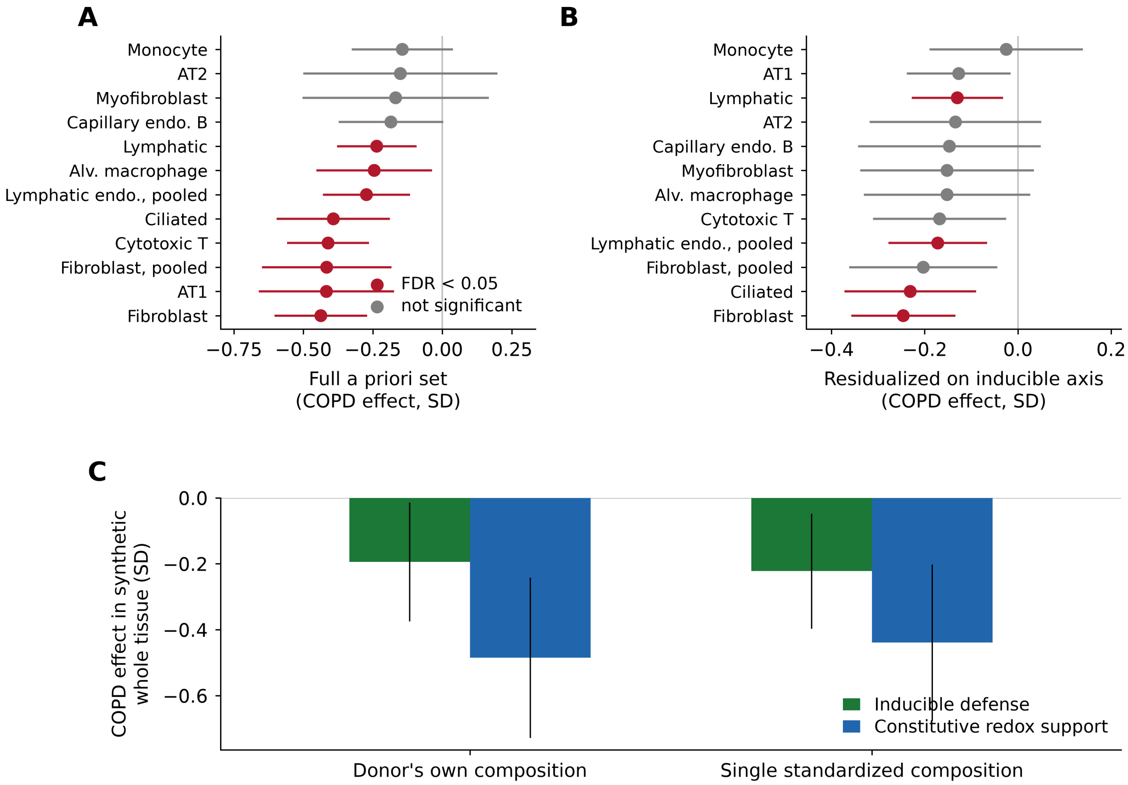


**Supplementary Figure S1.** The disproportionate constitutive deficit holds under alternative program definitions and cellular composition. (A) Constitutive-minus-inducible contrast by population using the complete a priori constitutive set with no genes removed, as donor-level estimates with 95% CI. Red marks Benjamini-Hochberg FDR below 0.05 and gray marks a non-significant estimate, as keyed on the panel. (B) The same difference using a constitutive score residualized against the inducible program, with the same color key. (C) Synthetic whole tissue recombined at each donor's observed composition and at a standardized reference composition, showing that composition within the discovery resource does not account for the cell-resolved effect. Both programs decline in synthetic tissue, with the larger loss in constitutive support.

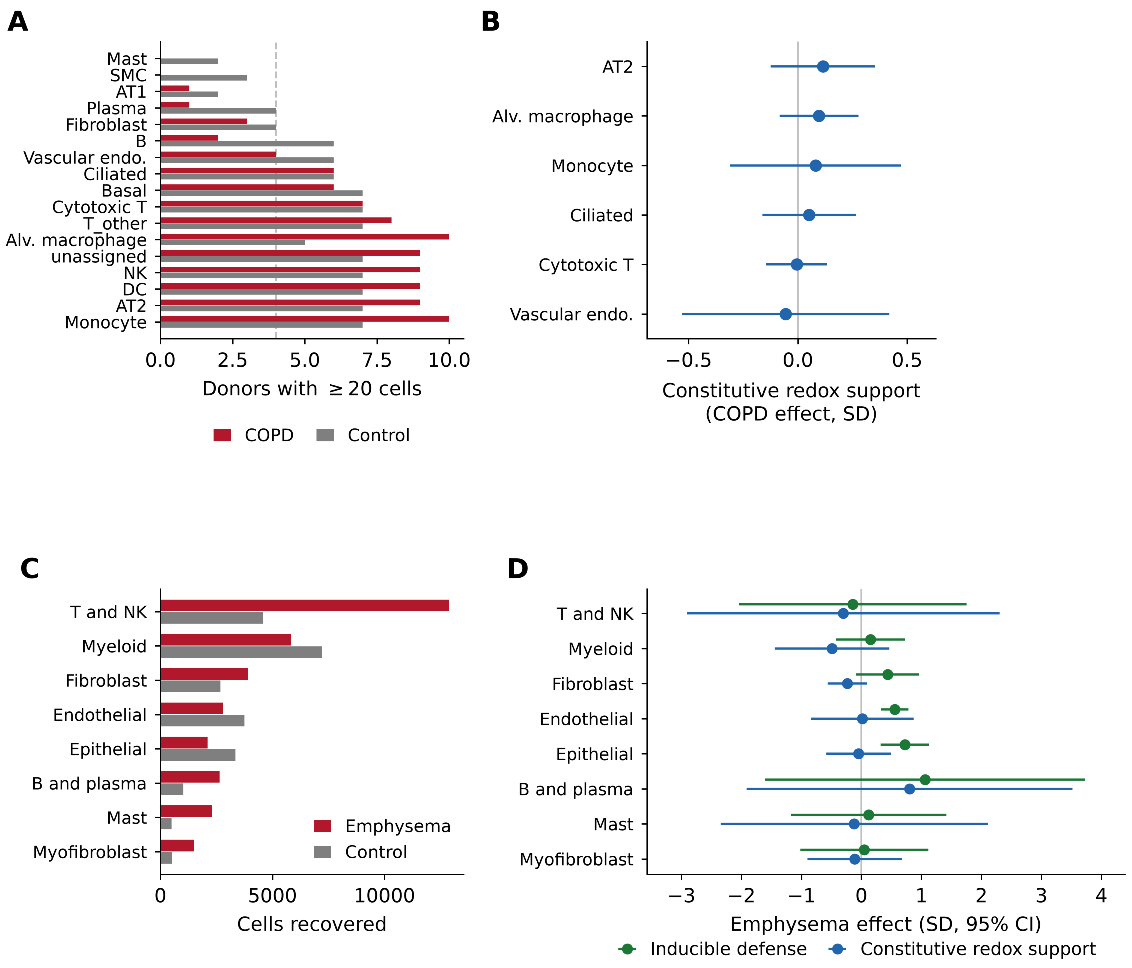


**Supplementary Figure S2.** What the independent cohorts can and cannot test. (A) Donor and cell coverage in the independent single-cell COPD cohort, showing which populations reach the donor depth required for testing. (B) Constitutive redox-support effect in the populations that do. (C) Cell recovery by population in the emphysema single-cell cohort. (D) Emphysema effect on each program in that cohort, with 95% CI. Neither cohort recovers fibroblasts and lymphatic endothelium at discovery depth, so these panels bound the replication rather than providing it.

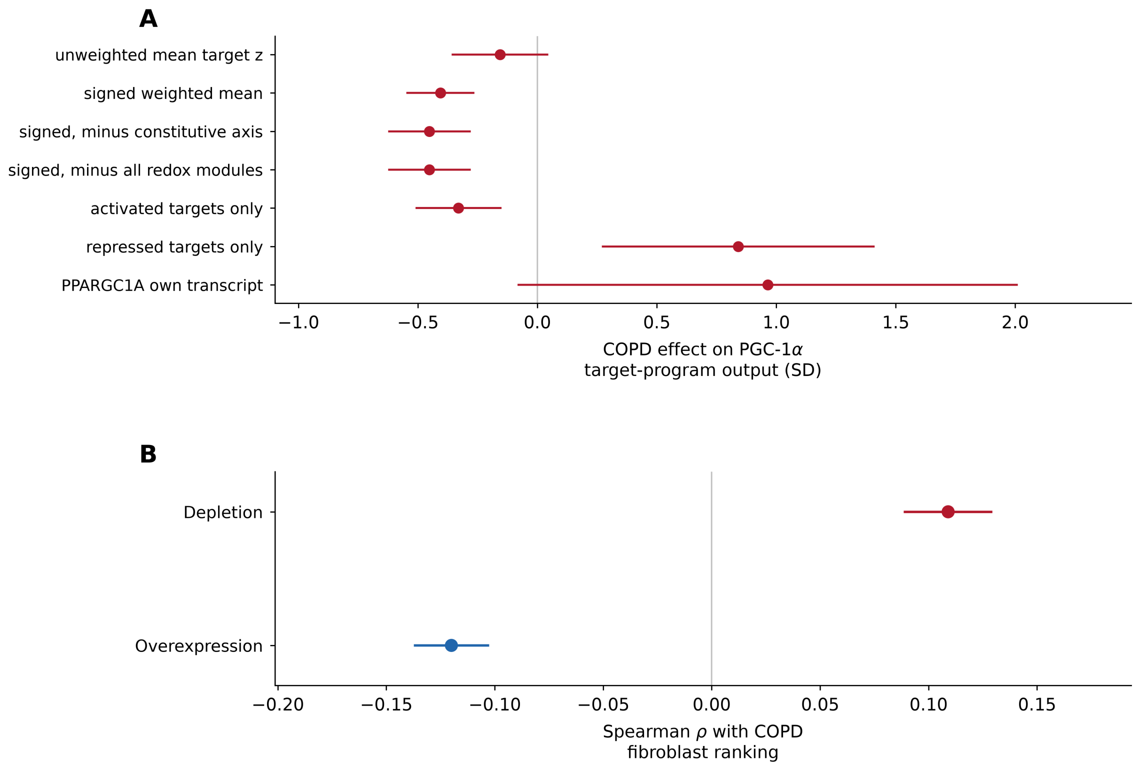


**Supplementary Figure S3.** Robustness of the PGC-1α target-program result. (A) COPD effect on inferred PGC-1α output in fibroblasts under seven estimator variants, all computed from the CollecTRI regulon, comprising unweighted and signed weighted target means, the signed mean after removing constitutive-support-axis genes and after removing all prespecified redox genes, activated targets only, repressed targets only, and *PPARGC1A* own transcript, with 95% CI. The five activity estimators are all negative. The repressed-target estimator is positive, which is the same biological direction because targets that PGC-1α represses rise when its output falls, and the transcript estimator is positive because it measures transcript abundance rather than target-program output. (B) Two independent PGC-1 perturbation signatures scored against the COPD fibroblast ranking. Neither was performed in lung fibroblasts.

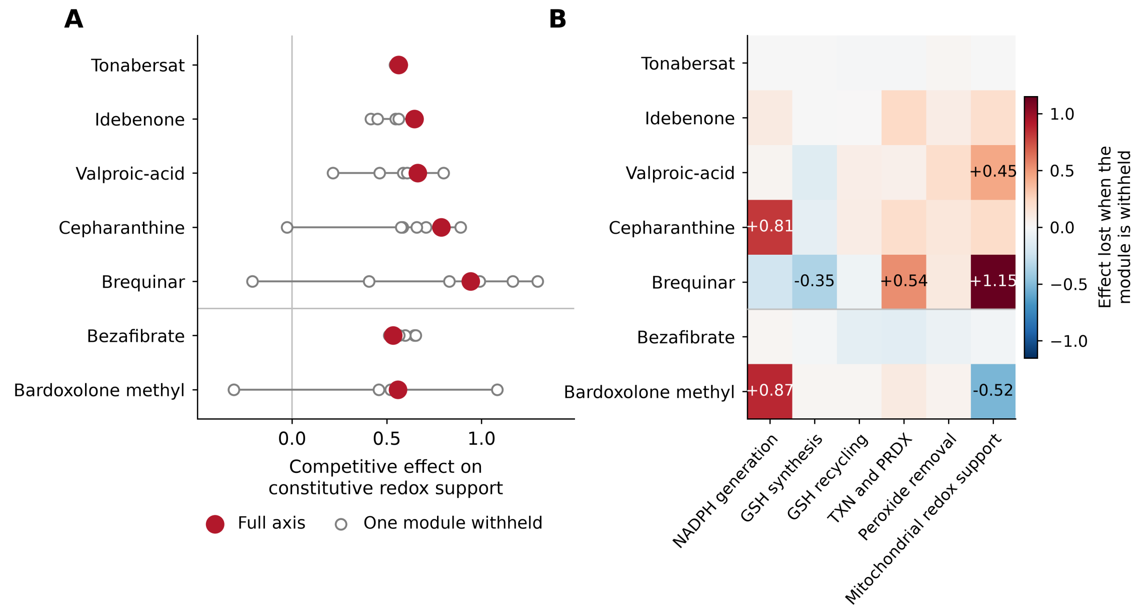


**Supplementary Figure S4.** Biochemical-arm dependence of the selected pharmacologic probes and the two named comparators. (A) Competitive effect on the constitutive program for each compound under the full program and under six recomputations that each withhold one biochemical module, with the five prespecified screen-derived probes above the rule and the two comparators below it. (B) The effect lost when each module is withheld, where positive values mark dependence on that module. Cepharanthine and bardoxolone methyl lose most of their effect when NADPH generation is withheld, brequinar depends most on mitochondrial redox support, and tonabersat and bezafibrate lose almost nothing to any single module. The full program was rescored within the same matched analysis set, producing minor differences from Figure 5. Arm dependence indicates which genes carry the score and is not by itself a mechanism.

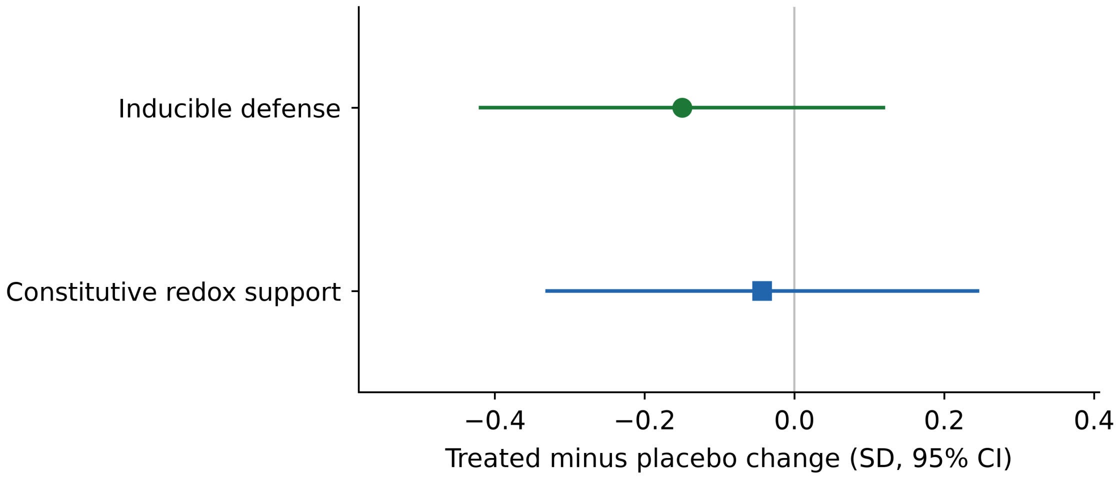


**Supplementary Figure S5.** Absolute treatment effects in the randomized nicotinamide riboside trial in COPD, shown as treated-minus-placebo changes in each standardized program score with 95% confidence intervals and on the trial's own absolute-effect scale rather than the competitive matched-null scale used in Figure 4B. Neither program reaches significance against zero on this test, while the inducible program is strongly displaced relative to expression-matched null gene sets in Figure 4B, so the two analyses answer different questions. Trial data, Dryad doi:10.5061/dryad.d2547d8b5.

Published evidence by experimental system

| Intervention | clinical COPD | COPD ex vivo | primary human lung | animal smoke | cell model | papers | trials | scored |
|---|---|---|---|---|---|---|---|---|
| N-acetylcysteine | ■ | ■ | ■ | ■ | □ | 21 | 15 | ● |
| carbocisteine | ■ | □ | □ | ■ | ■ | 16 | 0 | — |
| vitamin C / ascorbate | ■ | □ | □ | ■ | ■ | 15 | 4 | — |
| quercetin | ■ | ■ | ■ | ■ | ■ | 13 | 4 | ● |
| apocynin | ■ | □ | □ | ■ | ■ | 13 | 0 | ● |
| resveratrol | ■ | ■ | □ | ■ | ■ | 12 | 3 | ● |
| erdosteine | ■ | □ | □ | □ | □ | 10 | 4 | — |
| dimethyl fumarate | □ | □ | ■ | ■ | ■ | 9 | 0 | ● |
| curcumin | ■ | ■ | □ | ■ | □ | 8 | 3 | ● |
| bardoxolone methyl | □ | □ | ■ | ■ | ■ | 8 | 0 | ● |
| sulforaphane | □ | ■ | ■ | ■ | ■ | 6 | 2 | ● |
| vitamin E / alpha-tocopherol | ■ | □ | □ | □ | □ | 6 | 2 | — |
| MitoQ | ■ | □ | □ | ■ | □ | 5 | 2 | — |
| CDDO-imidazolide | □ | □ | ■ | ■ | ■ | 5 | 0 | — |
| cysteine | ■ | □ | □ | ■ | □ | 5 | 0 | — |
| glutathione | ■ | □ | □ | ■ | ■ | 3 | 0 | — |
| selenium | ■ | □ | □ | □ | □ | 2 | 2 | — |
| diphenyleneiodonium | □ | □ | □ | ■ | □ | 2 | 0 | — |
| genistein | □ | ■ | ■ | □ | □ | 2 | 0 | ● |
| metformin | □ | □ | □ | ■ | □ | 2 | 0 | ● |
| nicotinamide riboside | □ | □ | □ | ■ | □ | 1 | 1 | — |
| MitoTEMPO | □ | □ | □ | ■ | □ | 1 | 0 | — |
| melatonin | □ | □ | □ | ■ | □ | 1 | 0 | ● |
| pioglitazone | □ | □ | □ | ■ | □ | 1 | 0 | ● |
| procysteine | □ | □ | □ | ■ | □ | 1 | 0 | — |
| omaveloxolone | □ | □ | □ | □ | □ | 0 | 0 | — |
| bezafibrate | □ | □ | □ | □ | □ | 0 | 0 | ● |
| cepharanthine | □ | □ | □ | □ | □ | 0 | 0 | ● |

14 of 28 interventions had a compatible chemical signature and could be scored. The remainder are retained here with their evidence recorded and no effect assigned.

■ prior evidence   □ none identified   ● scored in this study   — no compatible signature

**Supplementary Figure S6.** The complete prior-evidence base for every redox-directed intervention returned by the prespecified search, ordered by weight of published evidence. Filled boxes mark published testing of that intervention in the named experimental system and open boxes none identified in this search, with the number of qualifying papers and registered trials given at right. The final column marks the interventions that had a compatible chemical signature and could be scored against the transcriptional programs. Several of the most heavily studied agents in COPD, including carbocisteine, vitamin C and erdosteine, have no compatible signature, so the transcriptional comparison in Figure 5 cannot reach them. Main Figure 5D displays a curated ten-row subset of this inventory, the five named comparators discussed in the Results together with five prespecified screen-derived probes, so the complete evidence for every intervention is given here rather than duplicated there. Evidence categories and inventory construction are in Methods, and the underlying records are Supplementary Table S11.
